# Bounds on the Ultrasensitivity of Biochemical Reaction Cascades

**DOI:** 10.1101/2023.02.28.529800

**Authors:** Marcello Pajoh-Casco, Abishek Vinujudson, German Enciso

## Abstract

The ultrasensitivity of a dose response function can be quantifiably defined using the generalized Hill coefficient of the function. Our group examined an upper bound for the Hill coefficient of the composition of two functions, namely the product of their individual Hill coefficients. We proved that this upper bound holds for compositions of Hill functions, and that there are instances of counterexamples that exist for more general sigmoidal functions. Additionally, we tested computationally other types of sigmoidal functions, such as the logistic and inverse trigonometric functions, and we provided evidence that in these cases the inequality also holds. We show that in large generality there is a limit to how ultrasensitive the composition of two functions can be, which has applications to understanding signaling cascades in biochemical reactions.

## 1 Introduction

The human body is a complex system with a large number of cell types working together to carry out different tasks. In some situations, cells need to be decisive in the sense of ignoring a low level of stimulus, while leading to a significant response when given a larger stimulus. For example consider the case of a wound, in which the skin breaks and bleeds. The surrounding cells immediately send signals to other cells in the skin, essentially telling them to divide quickly. This signal is sent in the form of a molecule called epidermal growth factor (EGF) which floats in the neighborhood of the wound, and which binds to a membrane bound receptor called epidermal growth factor receptor (EGFR) [2]. Cells that receive sufficient EGF binding to their receptors will begin to quickly divide. On the other hand, this is a very sensitive system since unwanted replication can lead to cancer in contexts other than wound healing. In order to prevent such unwanted cell division, the EGFR dose response is such that a small amount of EGF leads to no response, while a slightly larger input EGF concentration leads to a robust response. This behavior is called *ultrasensitivity*.

A highly ultrasensitive switch is similar to a bell: when you press the button sufficiently it rings, but it shows no response for a light press [4]. In the context of EGFR signaling, the molecules downstream of this receptor are modified by phosphorylation [4], the transfer of phosphate molecules mediated by an enzyme. Phosphorylation takes place sequentially over a series of molecules, each molecule phosphorylating (and thereby activating) the next. This particular cascade is known as the Mitogen-Activated Protein Kinase cascade (MAPK), and it is model signaling pathway in the study of cellular communication [10]. It is believed that the combination of these multiple steps is in large measure what allows the larger ultrasensitivity of the overall response.

In this paper, we study how connecting a cascade of multiple small reactions with moderate ultrasensitivity can result in a single cascade with significantly larger ultrasensitivity. Specifically, we work to establish an upper bound to the extent of ultrasensitivity in a cascade, in terms of the ultrasensitivity of the individual cascade steps. In the context of the MAPK cascade example, each of the three steps constitutes a smaller set of reactions with its own input-output response, and the overall dose response of the system can be broadly understood as the composition of each of these functions. We therefore ask how ultrasensitive can be the composition of moderately ultrasensitive functions.

In first instance we use Hill functions to describe the input-output behavior of each cascade step. This function can also be used to quantitatively measure the ultrasensitivity of a response through the so-called Hill coefficient, a component of the function referred to as *n* in the formula below and Figure 1a): [16]

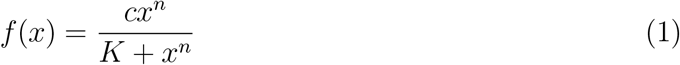

In this case *x* is the input concentration of ligand [14], and *c* describes the saturation value of the function for large *x*. The constant *K* modulates the function horizontally, such that when *x^n^* = *K* the response is 50% of the maximal output.

**Figure 1:**
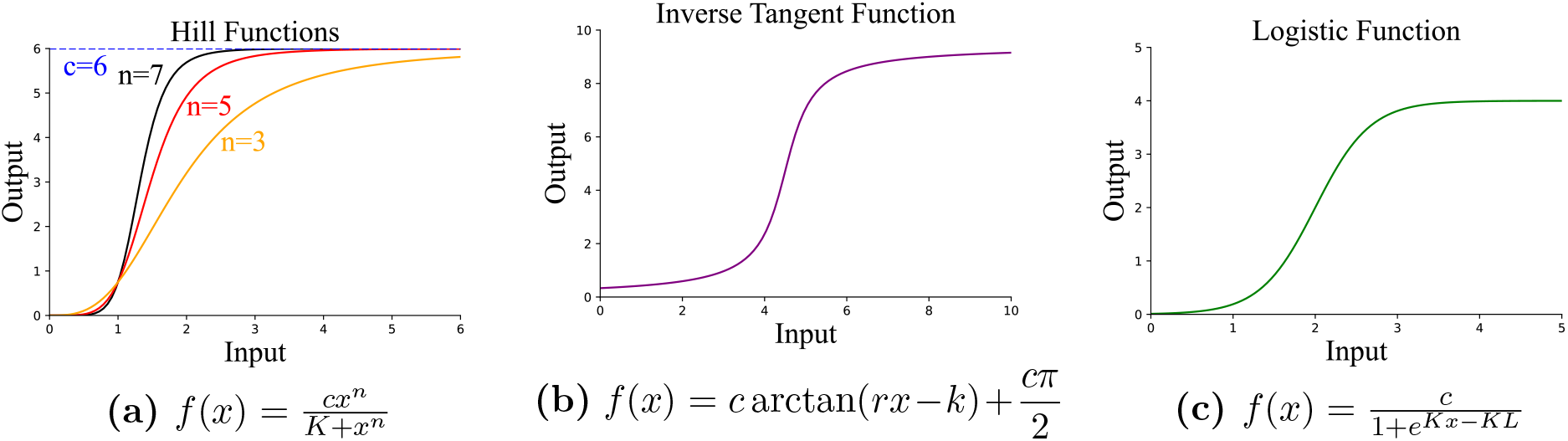
**a)** Hill function graphs, each with a different value of *n* to illustrate how the Hill coefficient affects the properties of each function. All functions on the graph were evaluated at *c* = 6 and *K* = 7. **b)** The graph of an inverse tangent function evaluated at *c* = 3, *r* = 2, and *K* = 9. **c)** The graph of a logistic function evaluated at *c* = 4, *K* = 3, and *L* = 2.

Given a positive, increasing, saturating function *f*(*x*), we can also define the values

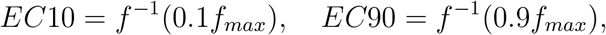

where *f_max_* is the saturation value of the function [1,7]. They are the effective concentrations of input that lead to 10% and 90% of the response, respectively. These values can also be used to find the Hill coefficient for non-Hill functions through the use of the formula derived by Goldbeter and Koshland in 1981 [5]:

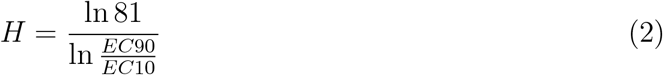

While the value of *H* can be calculated for any such so-called *sigmoidal* function, it also satisfies that *H* = *n* in the case of Hill functions. This generalized Hill coefficient is therefore helpful to quantify the ultrasensitivity of dose responses in larger generality.

The main conjecture we explore in this paper is inspired by work proposed by James Ferrell and colleagues. We propose and prove that the Hill coefficient of the composition of two Hill functions *f*(*x*), *g*(*x*) satisfies the formula

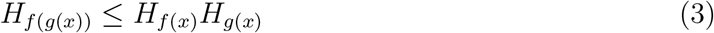

The value of this hypothesis, which we call the Ferrell inequality, is that it provides an upper limit for how ultrasensitive a cascade can be, as a function of the individual steps. We establish here the inequality for two arbitrary Hill functions. We explore computationally the generalization of the inequality to three-step Hill function cascades, as well as for two other families of sigmoidal functions, namely inverse trigonometric functions and logistic functions. Finally, we show that the inequality is not true for any two arbitrary sigmoidal functions, by producing a simple counterexample.

In the work [8], Huang and Ferrell show a similar inequality in terms of the sensitivity of a dose response, rather than the Hill coefficient. The *sensitivity* of a function *f*(*x*) is defined as *S*(*x*) = *f′*(*x*)*x/f*(*x*). One can consider sensitivity as the percent change in the output based on a percent change in the input *x*. For example, a sensitivity of 3 means that an increase in the parameter x of 1% will result in an increase of the output by 3%. Huang and Ferrell pointed out a result for compositions of multiple functions, namely that

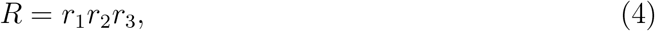

where R is the sensitivity of the cascade, and *r*_1_, *r*_2_, and *r*_3_ are the sensitivities of each level of the cascade. The proof of this result follows immediately from using the chain rule in the definition of the composition.

The sensitivity of a function is also used as a measure of ultrasensitivity, but it has some caveats. For instance, the sensitivity of a Hill function has the somewhat unintuitive property that it is highest at *x* = 0. The Hill coefficient, especially for Hill functions, is a widely followed measure of ultrasensitivity, and it is a natural question to explore this inequality in that context.

## 2 Results

### 2.1 Logarithmic Sampling

In order to ensure that the trials produced by our algorithm are properly randomized, we utilized logarithmic sampling. Suppose a lower bound and an upper bound are given for the sample, 0 < *a* < *b*. Since *b* might be orders of magnitude larger than *a*, we want to ensure that the samples are selected from each order of magnitude with similar likelihood. To do this we choose a number *x* between log_10_ *a* and log_10_ *b*, selected using a uniform distribution. The output of the logarithmic sampling algorithm is 10^*x*^, which must lie between *a* and *b*.

For instance, suppose that *a* = 1 and *b* = 1000 describe a plausible range for the parameter value of a given biochemical constant. If a number is sampled uniformly from 1 to 1000, most of the numbers sampled will be larger than 100. Using logarithmic sampling, one would choose a number *x* from 0 to 3, and the sampled number would be 10^*x*^. In this way, the sample has the same likelihood of being on the intervals [1,10], [10,100], and [100,1000].

### 2.2 Hill Function Database

We chose to computationally analyze Ferrell’s hypothesis using Hill functions in order to gain confidence that an analytical proof can be pursued. Two Hill functions *f*(*x*) and *g*(*x*) are considered, by randomizing each of their parameters using logarithmic sampling.

The Hill coefficient of *f*(*x*) and *g*(*x*) is simply given by the parameter *n* in each function. For the composition, the generalized Hill coefficient is calculated by inverting each function to calculate *EC*10 and *EC*90. Should *EC*10 be negative, the full set of parameters is randomized again to ensure that the generalized Hill coefficient H is well defined.

The ultrasensitivity of the composition function, *f*(*g*(*x*)), was then compared with the product of the ultrasensitivity of *f*(*x*) and *g*(*x*). The algorithm would then output a “yes” or “no” in order to signify if the values were consistent with our hypothesis. There were over 5000 trials performed. A portion of the database created by the algorithm can be seen below for illustrative purposes. Figure 1a) shows three Hill functions and demonstrates how changes in the Hill coefficient can modify its characteristics.

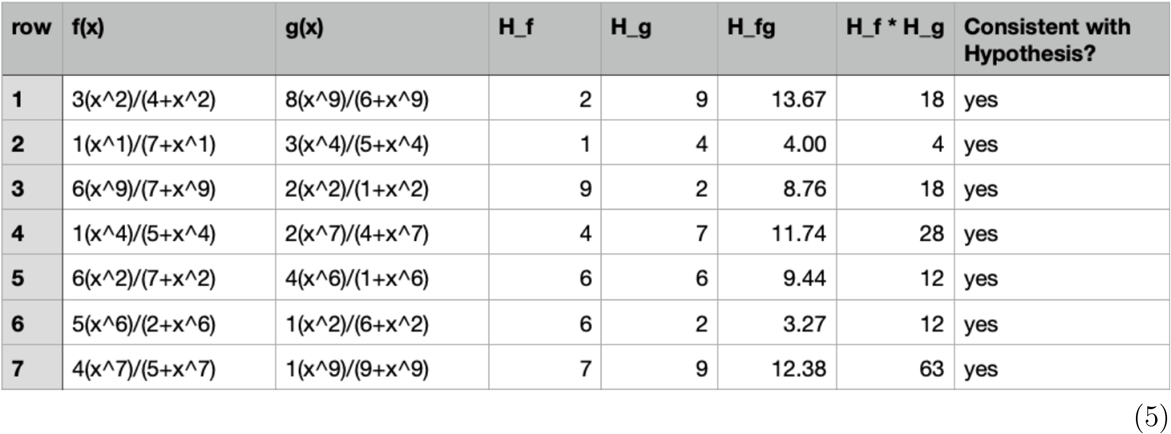

Every case that arose from the 5000 randomized trails in our database was consistent with the hypothesis, providing significant evidence that an analytical proof can be pursued in the case of two Hill functions.

### 2.3 Inverse Tangent Function Database

While constructing and analyzing the Hill function database, we recognized that we could computationally test our hypothesis with additional sigmoidal functions. We discuss our results using the inverse tangent function,

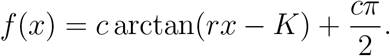

An example of the inverse tangent function can be found in Figure 1b. The parameters *c*, *r*, and *K* were randomized within biological parameters for the various trial cases using the logarithmic sampling method described in Section 2.1. We analyzed over 5000 trial cases for our hypothesis using inverse tangent functions. A portion of the database created by our algorithm can be seen below for illustrative purposes.

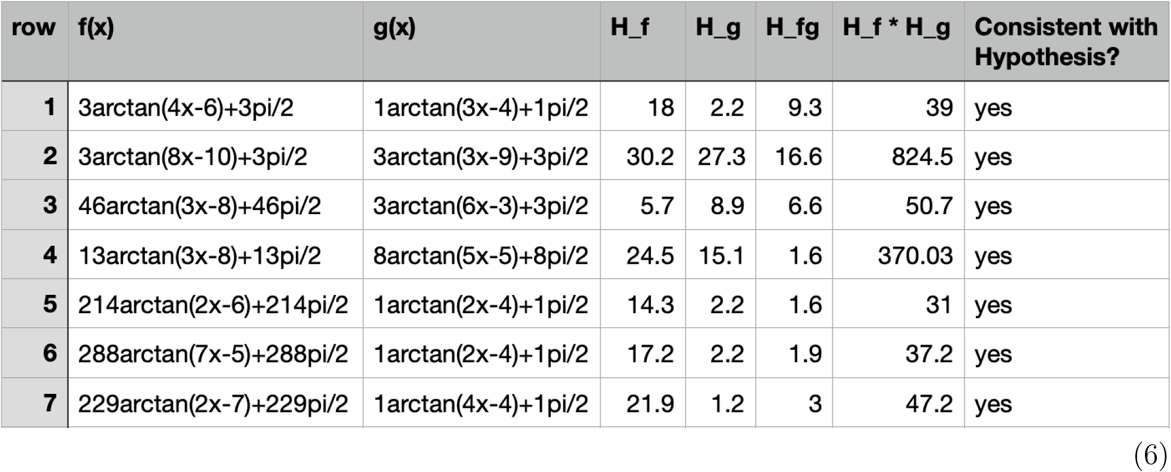

Over 98% of the cases that arose from the 5000 randomized trails in our database were consistent with our hypothesis. These results indicated that our hypothesis could be proven true not just in the case Hill functions, but other sigmoidal functions used to describe ultrasensitive responses as well. The remaining 2% of cases could be due to a failure of the inequality itself, or it could be due to numerical issues when calculating the Hill coefficients.

### 2.4 Logistic Function Database

The logistic function is another type of sigmoidal function that we chose to test our hypothesis computationally. We defined the logistic function as

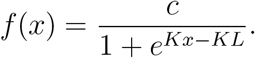

An example of the logistic function can be found in Figure 1c. The parameters *c*, *L*, and *K* were randomized within biological parameters for the various trial cases using the logarithmic sampling method described in Section 2.1. We analyzed over 5000 trial cases for our hypothesis using logistic functions. A portion of the database created by our algorithm can be seen below for illustrative purposes.

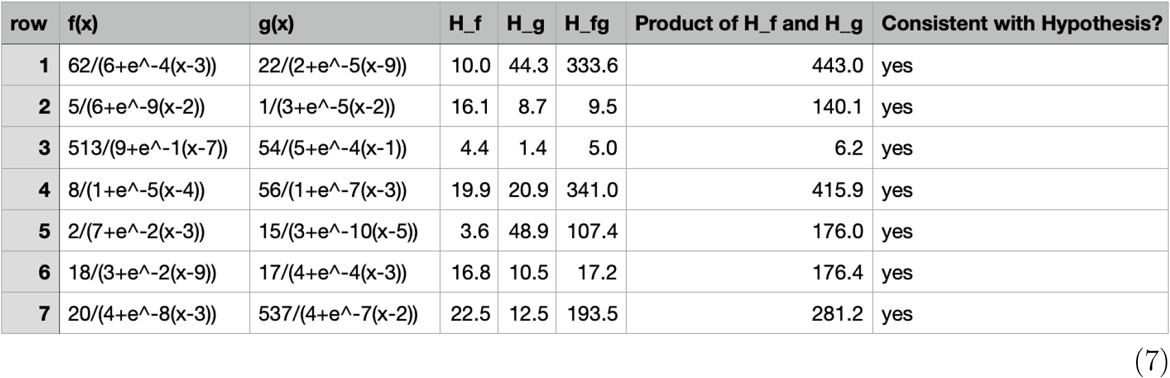

Every case that arose from the 5000 randomized trials in our database was consistent with our hypothesis. These results highlighted yet another instance in which our hypothesis could be proven true by other sigmoidal functions used to describe ultrasensitive responses.

### 2.5 Proving the Ferrell Inequality for Hill Functions

In this section we provide a rigorous mathematical proof for the Ferrell inequality in the case of two Hill functions. We start this proof by establishing the notation for the three functions *f*(*x*), *g*(*x*), and *h*(*x*) = *f*(*g*(*x*)),

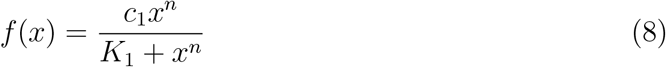

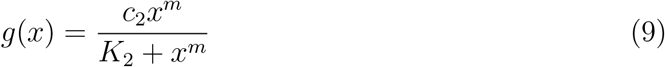

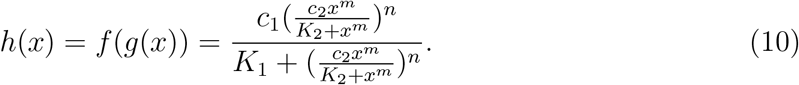

Recall that we want to prove

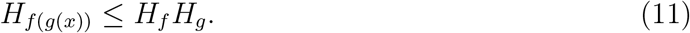

First, we assume without loss of generality that *c*_1_ = 1 and *c*_2_ = 1. Suppose that the result is true in that special case. In the general problem, *c*_1_ only re-scales *f*(*x*) and *h*(*x*) vertically. It does not affect the Hill coefficients involved.

Similarly for *c*_2_,

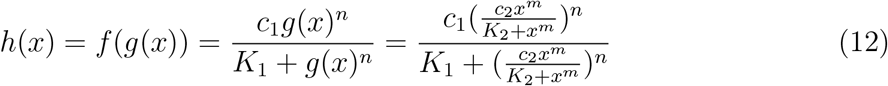

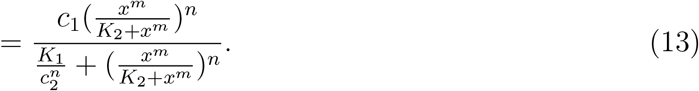

Notice that the composition function *h*(*x*) is identical to that of the case *c*_2_ = 1, in which the coefficient *K*_1_ has been replaced by 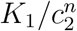. In this way, one can assume that *c*_1_ = 1 as well as *c*_2_ = 1 without loss of generality, and we do this in the rest of the proof.

In order to prove the inequality, we first have to determine the ultrasensitivity, *H*, by calculating the Hill coefficient for each function. Since *f*(*x*) and *g*(*x*) are Hill functions, we already know that *H*_*f*(*x*)_ = *n* and *H*_*g*(*x*)_ = *m*. However, determining the Hill coefficient for the composition function *f*(*g*(*x*)) will prove to be more challenging since it does not follow a form which allows us to directly determine its ultrasensitivity using the Hill coefficient as we did with *f*(*x*) and *g*(*x*). Instead, we will have to use equation (2), 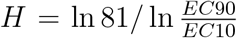. We will first find the maximum *y* value of the composition function, then use the inverse of the composition function to solve for the EC10 and EC90 values. We find the ratio of the EC90 to EC10 values, and then insert that ratio into equation (2).

For the composition of two Hill functions, the exterior function evaluated at the saturation point of the interior function will produce the maximum *y* value, or *y_max_* = *f*(*c*_2_). Recall we assumed *c*_2_ = 1 without loss of generality. Evaluating *f*(*c*_2_) results in

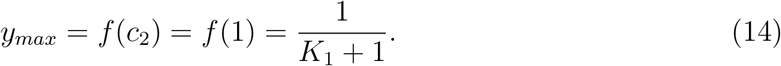

Now that we have the maximum *y* value of the composition, we can solve for the EC10 and EC90 values. Starting with the EC90, we need to solve for *x* in the equation

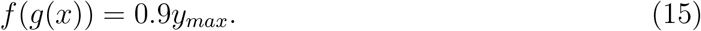

Evaluating the inverse functions on both sides we get

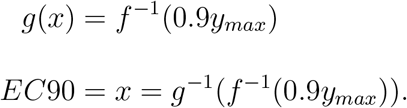

We can take the following steps to derive the formula for the inverse of a Hill function:

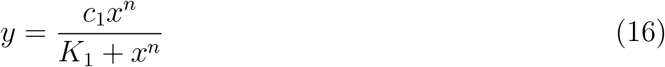

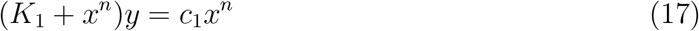

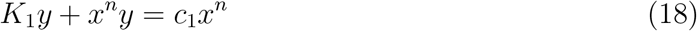

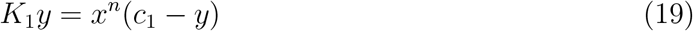

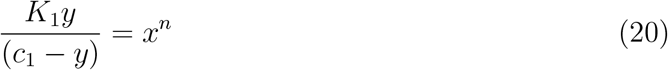

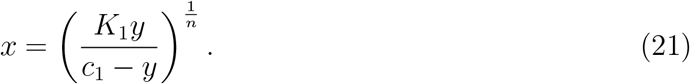

The inverse formula for *g*(*x*) is analogous. Now that we have the formula for the inverse of a Hill function, we can apply it to the formula *EC*90 = *g*^−1^(*f*^−1^(0.9*y_max_*)) as we described previously. This yields

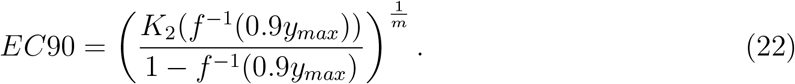

We can follow the same process as above to solve for the EC10 value derived from the equation *EC*10 = *g*^−1^(*f*^−1^(0.1*y_max_*)), which results in

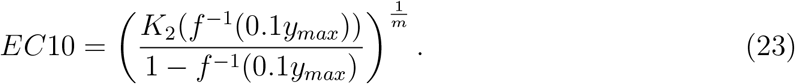

In order to further simplify the EC90 and EC10 expressions, we can write *f*^−1^(0.1*y_max_*) and *f*^−1^(0.9*y_max_*) in terms of *K*_1_ and n, and relabel them as *α* and *β*, respectively, for clarity when being referenced

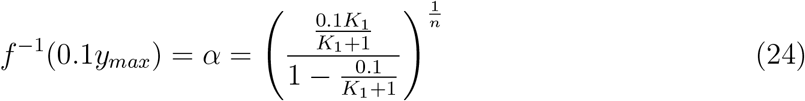

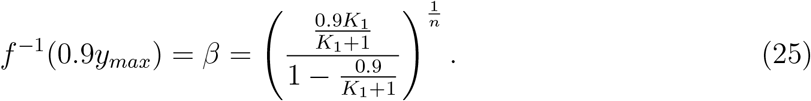

Let us take the time to further simplify *α* and *β*. For *α*, we can start the simplification process by getting a common denominator for the expression in the denominator of *α*,

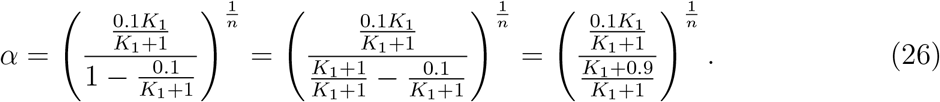

Cancelling fractions and further simplifying we get

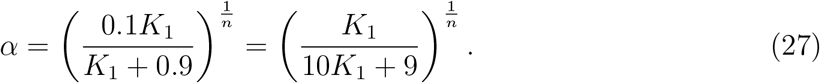

Following the same process we did for *α* will allow us to reduce *β* to

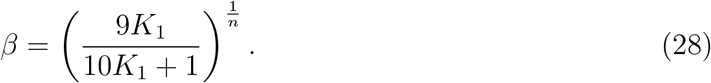

Notice that 0 < *α* < *β* 1, which will be important below.

Now that we have solved for both the EC10 and EC90 values and simplified them in terms of *α* and *β*, we can form the EC90 to EC10 ratio,

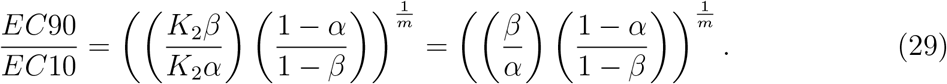

We can then take this expression and insert it into equation (2) in order to determine the ultrasensitivity of the composition function,

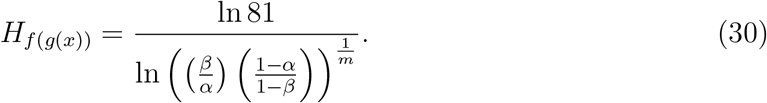

Now that we have determined the ultrasensitivity for all three functions, *f*(*x*), *g*(*x*), and *f*(*g*(*x*)), we can insert them into our hypothesis, the Ferrell inequality:

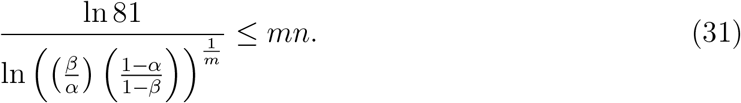

In order to simplify this inequality, we can multiply both sides by the natural logarithm expression and apply the properties of logarithms to cancel out *m*,

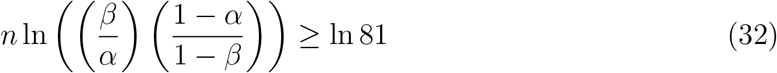

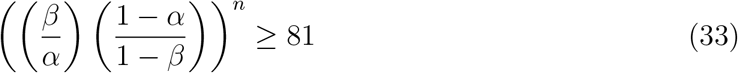

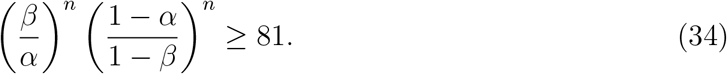

In the inequalities above we used the fact that *β/α* > 1 and (1 – *α*)/(1 – *β*) > 1 so that the logarithm in the numerator is positive. Notice also that this expression, which is equivalent to the Hill inequality, only depends at this point on the parameters *n* and *K*_1_. Since *α* > 0 and 1 – *β* > 0, this expression is equivalent to

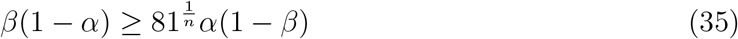

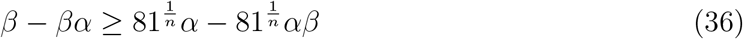

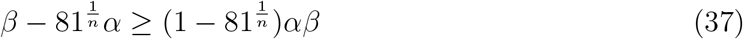

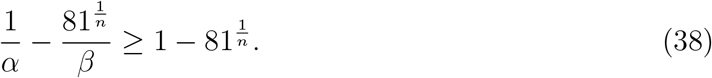

In order to prove this expression, let us first consider the case for *n* = 2:

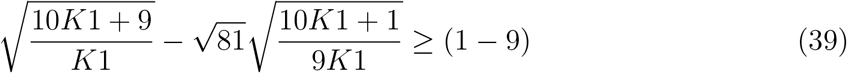

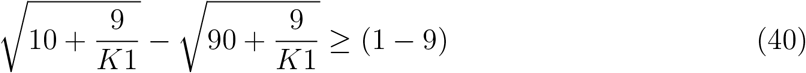

We multiply by the conjugate

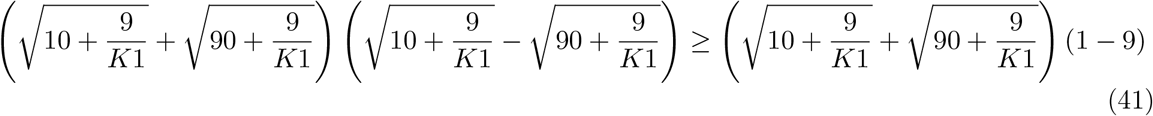

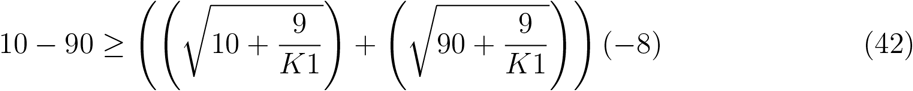

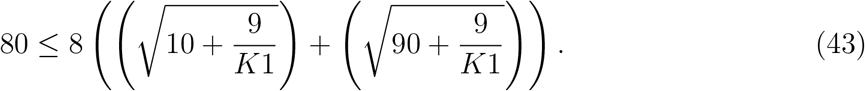

Since the right hand side is decreasing as a function of *K*_1_, it is sufficient to evaluate this inequality as *K*_1_ approaches ∞,

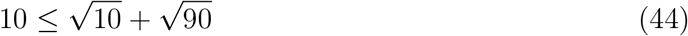

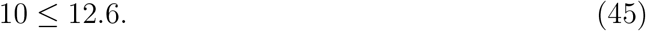

Now let us consider the case where *n* is arbitrary. We start by applying the following algebraic identity:

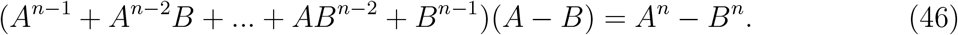

We define Λ as the generalized conjugate expression from the case *n* = 2,

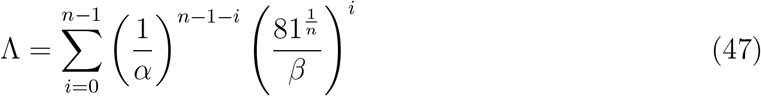

By multiplying the inequality on both sides with Λ we get

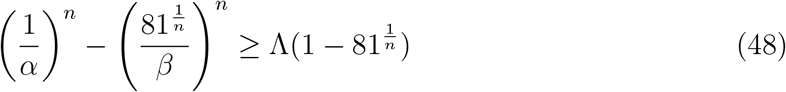

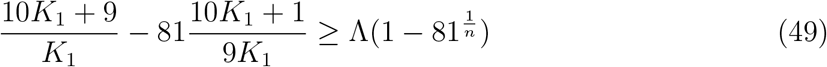

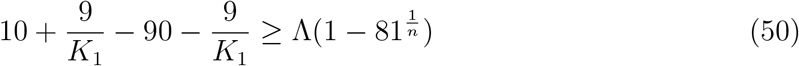

We can now cancel out 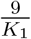 from the left hand side and multiply the equation by −1, to get

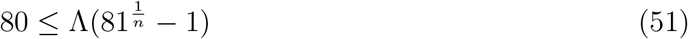

Recalling the expression Λ, we calculate

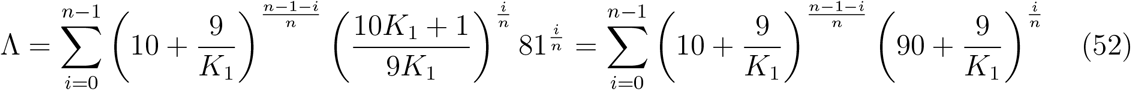

Since Λ is a decreasing function of *K*_1_, to prove equation 23 true, it is sufficient to prove it in the case where *K*_1_ approaches ∞:

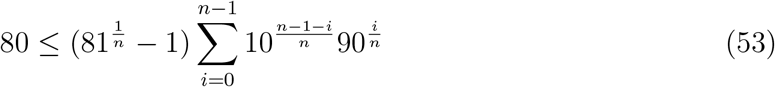

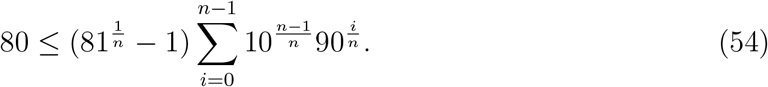

In order to evaluate the series recall that

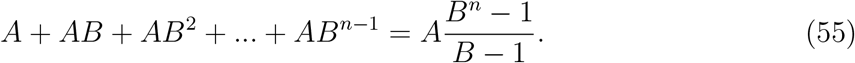

If we evaluate the last inequality by applying the above identity, where 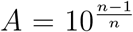 and 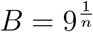, we get

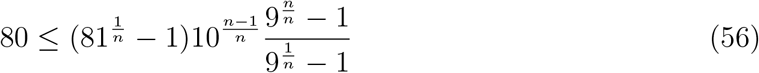

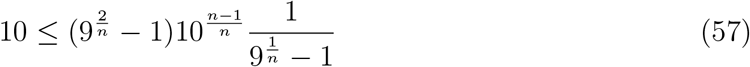

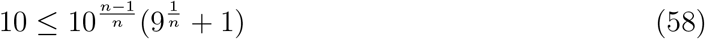

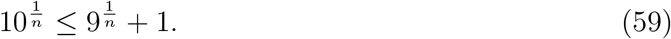

This expression is equivalent to

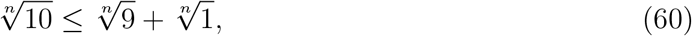

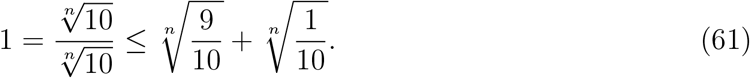

Now, notice that

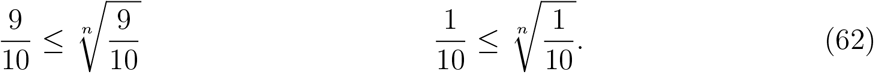

The result follows by adding both of these inequalities,

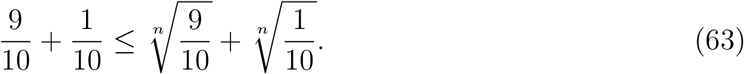

In this way, we have proven the Ferrell inequality for all Hill functions.

### 2.6 Counterexample to the Ferrell Inequality

As seen through our results thus far, we have analytically proven that the Ferrell inequality holds for all Hill functions. There is also significant evidence through our computational work that the inequality holds for other sigmoidal functions. However, there are counterexamples for certain sigmoidal functions. Recall our hypothesis,

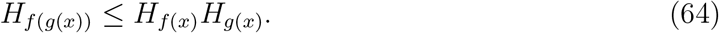

Define the following functions:

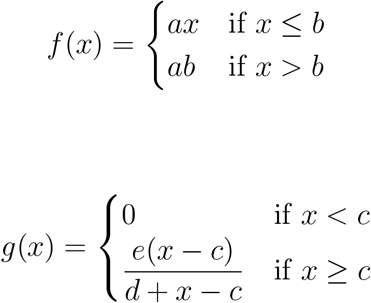

where *a*, *b*, *c*, *d*, and e are all parameters. The function *f*(*x*) is a diagonal function that saturates, while the function *g*(*x*) is a Hill function with *n* = 1 that has been shifted to the right.

We can calculate the 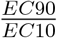 ratio and use it to solve for the Hill coefficient of all three functions. In the case of *f*(*x*) we know that the maximum y-value is *b*. We can use the properties of piecewise functions to calculate the the ratio. By the definition of f(x), we know that y is equal to x, so we can determine the EC10 and EC90 values by calculating 10% and 90% of the maximum,

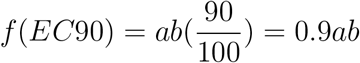

and:

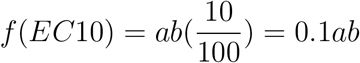

With this information, we use the *f*(*EC*90) and *f*(*EC*10) values to find the corresponding EC90 and EC10 values of the function, which are 0.9*a*^2^*b* and 0.1*a*^2^*b*. Then, by utilizing equation (2),

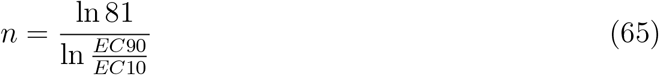

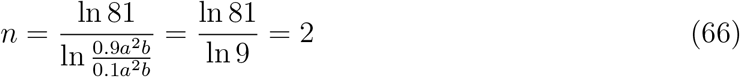

We know that the Hill coefficient of *f*(*x*) is 2 no matter what the values for a and b are. Similarly, we can determine the Hill coefficient of *g*(*x*). The maximum value for *g*(*x*) is e since as *x* becomes an infinitely large number, *c* and *d* in (*x* – *c*) and (*x* + *d* – *c*) become negligible. The value e the becomes the maximum value since 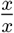 reduces to 1. We know that *g*(*EC*90) and *g*(*EC*10) are

We calculate the *EC*90 value as follows,

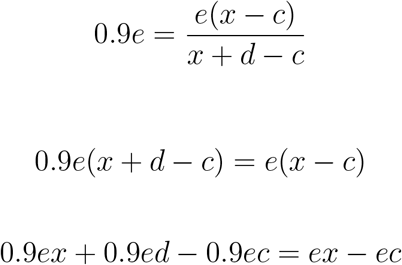

Simplifying the terms on each side yields

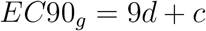

Similarly, we can calculate the *EC*10 value, 9x-9c = d

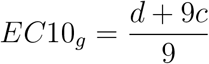

We can plug both of these values, the EC90 and EC10, into the Goldbeter and Koshland formula below:

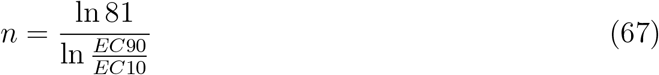

so:

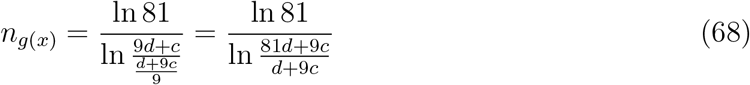

In order to calculate the Hill coefficient of the composition, notice that by construction the composition consists of truncating the function *g*(*x*) at the value *ab*. The composition has this maximum value and we have

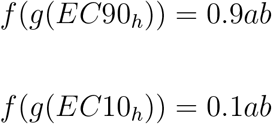

We can do a similar calculation as for the function *g*(*x*),

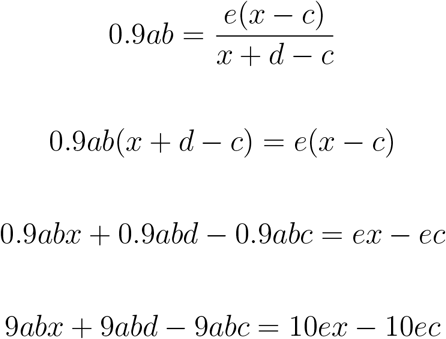

We can find the value for the EC90 of *f*(*g*(*x*)) by simplifying this equation and substituting *EC*90_*f*(*g*(*x*))_ for x:

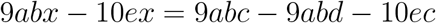

so:

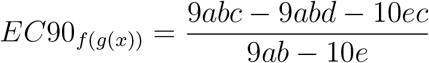

We can apply a similar method to find the EC10,

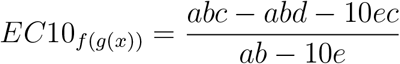

For example, let us use the parameters *a* = 1, *b* = 2, *c* = 3, *d* = 4 and *e* = 25. Then, by plugging in values to the EC10 and EC90 formulas we can go back to *g*(*x*),

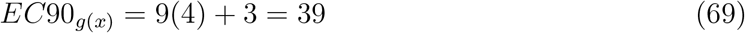

and the EC10 for *g*(*x*) is:

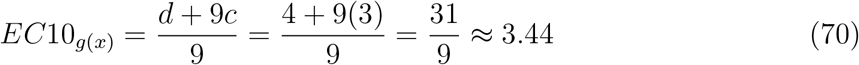

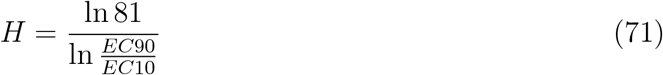

so:

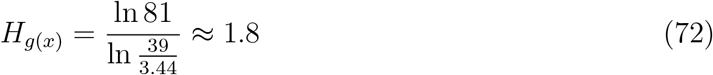

The product of *f*(*x*) and *g*(*x*) for this specific parameters is 3.62. We plug in the same values to find the *EC*90_*f*(*g*(*x*))_ and the *EC*10_*f*(*g*(*x*))_.

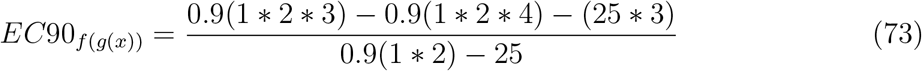

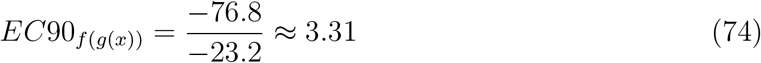

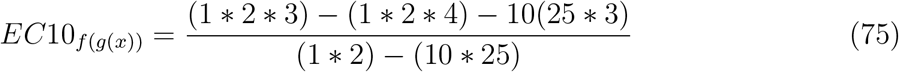

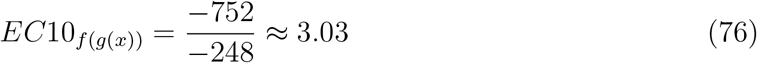

We calculate

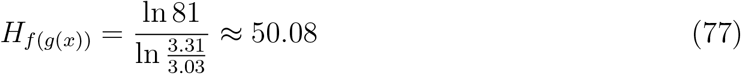

Notice how neither *f*(*x*) nor *g*(*x*) are particularly ultrasensitive. To reiterate, the Hill coefficients are 3.31 and 1.80, respectively. However, the Hill coefficient of the composition is 50.08, which is highly ultrasensitive. Figure 2a presents the graph of *f*(*x*) in red, Figure 2b presents the graph of *g*(*x*), Figure 2c presents the graph of *f*(*g*(*x*)),

**Figure 2:**
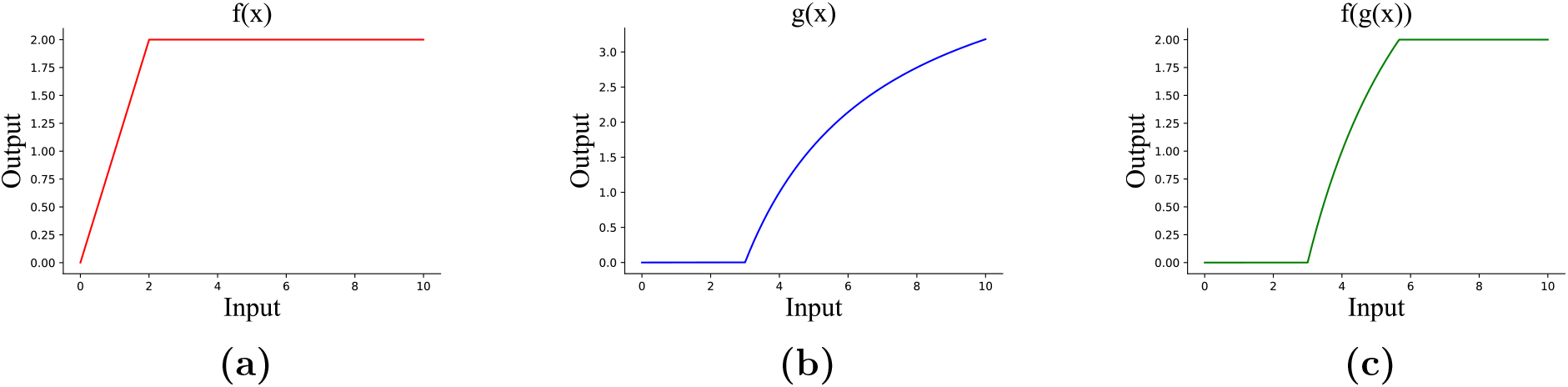
Counterexample to Ferrell’s Inequality. **a)** Graph of the function *f*(*x*), **b)** Graph of the function *g*(*x*), **c)** Graph of the function *fg*((*x*)) as defined in this section on the domain of 0 to 10.

Figure 2 shows how neither *f*(*x*) nor *g*(*x*) are particularly ultrasensitive. However, the graph for *f*(*g*(*x*)) increases rapidly and then flattens at a relatively close y-value, resulting in a large amount of change in the y-axis from a small amount of change in the x-axis.

### 2.7 Three Hill Function Analysis

In order to expand the horizons of our research, our team has analyzed the composition of three Hill functions. The reason for this is to see if our hypothesis will still hold true for more than just two Hill functions. Additionally, our research focuses on improving the ultrasensitive response of a cascade. If we are able to make those cascades more switch like by composing more functions together, it would be more effective. In order to establish this change, we had to slightly modify our hypothesis for this case to be as follows.

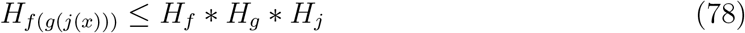

We have the third cascade to be denoted as *j*(*x*). Thus, our team generated a database of 5000 rows that found that in all 5000 cases the values were consistent with the hypothesis.

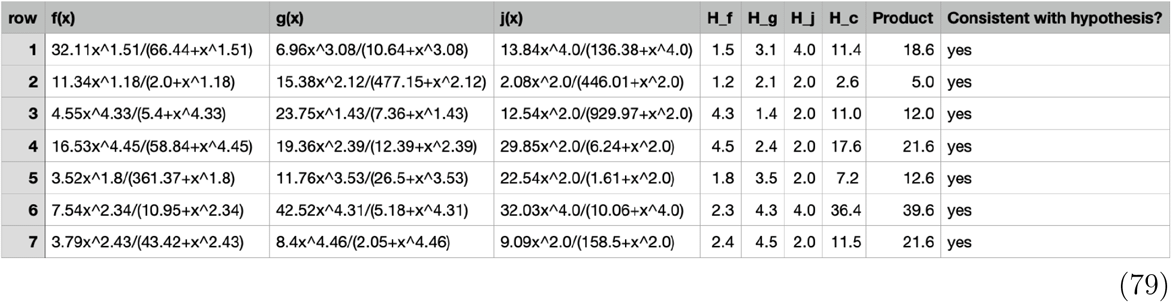

The high probability that our modified hypothesis is applicable to more than just one composition expands the range of our research where we can significantly improve the ultrasensitivity of a cascade by composing it with more than one other cascade.

## 3 Discussion

### 3.1 Ferrell Inequality

An analysis of the Ferrell inequality has shown us that it holds true for all Hill functions, but not all sigmoidal functions. We have shown that in large generality there is a limit to how ultrasensitive the composition of two functions can be. From these results, it is evident that when using Hill functions to model the ultrasensitivity of a biochemical system, that a function with low ultrasensitivity and high sensitivity cannot be combined in a clever way to produce a composition function with high ultrasensitivity. There must already be two functions with high ultrasensitivity to produce a composition function with high ultrasensitivity. It is also important to note, however, that sigmoidal functions other than the Hill function can be used to analyze and represent biological systems. Since Ferrell’s hypothesis does not hold true in all cases, we cannot predict with absolute certainty the characteristics of ultrasensitivity in these biological systems in the same way we can for just Hill function-based systems.

### 3.2 Databases

While Hill functions are useful as the “standard” sigmoidal function for our purposes, we also wanted to check to see if our hypothesis worked with other cases as well. Each row of data translates to a possible situation that could happen in a cell and by having lots of data we are able to confirm if our hypothesis works in a real environment. The database also poses future implications of our project because we could use it to find the appropriate dosage required for a certain treatment. For this part of our project we decided to expand our horizons into other functions such as the inverse tangent and logistic since they are also sigmoidal. We wanted to make sure that our hypothesis wasn’t limited to just Hill functions and that it worked for other sigmoidal functions. Part of our reasoning for the hypothesis is that in order to get a certain output, we must put in enough input so that even in non-ideal conditions, having enough input can allow for the chance for some output to occur. This also relates to another implication of our project in the long term as by expanding our range of functions in which our hypothesis works for, we can guarantee a higher reliability in its actual biological implications. Our goal in this paper is to analyze ways to make certain functions as “switch-like” as possible so by analyzing different functions and testing varying parameters, we can see which combination of functions will give us the most switch like response. Thus, by also incorporating different cases of function, out team hopes to see if our hypothesis can go beyond the current state of our project and if it is applicable in many other cases. Moreover, as we created a Hill function table, and were able to numerically prove the case for the composition of two Hill functions, we wanted to expand our project and were able to create a table for three Hill functions. For 5000 rows, the three Hill function database showed that the results were consistent with our hypothesis and that there was a very high chance that it could be proven mathematically.

### 3.3 Proof of Goldbeter-Koshland Result

In this short section we include a proof of the result by Goldbeter and Koshland, generalizing the definition of the Hill coefficient *n* to sigmoidal functions that may not be Hill functions. The only aspect left to show is that in the case of Hill functions, *H* = *n*. Suppose

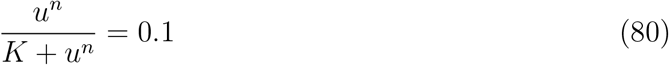

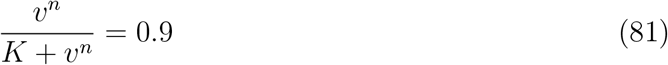

We can invert these two equations to produce the following

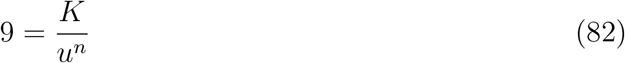

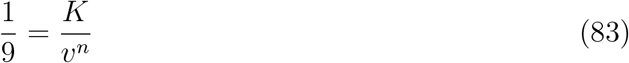

Then the first equation can be divided by the second, resulting in

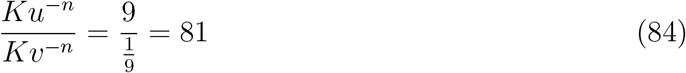

This equation can then be further reduced by taking the natural logarithm of both sides and solving for *n*, which produces the final result of

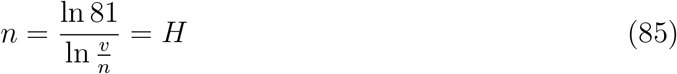

Once again, *n* is the value of the Hill coefficient, which we can use to quantify ultrasensitivity. The reason that EC10 and EC90 are used to calculate the Hill coefficient in the equation above instead of other possible values is because they provide information about the beginning and the end of the response curve.

